# INPP5E controls ciliary localization of phospholipids and odor response kinetics in a mouse model of Joubert syndrome

**DOI:** 10.1101/451344

**Authors:** Kirill Ukhanov, Cedric Uytingco, Warren Green, Lian Zhang, Stephane Schurmans, Jeffrey R. Martens

## Abstract

Ciliopathies manifested in part by a dysfunction of several phosphoinositide 5’phosphatases constitute Lowes, Dent disease 2 and Joubert syndromes through critical involvement of properly functioning primary cilia (PC). We showed that deletion of INPP5E under the control of OMP-Cre in mature mouse olfactory sensory neurons (OSNs) led to a dramatic redistribution of PI(4,5)P2 (PIP2) in cilia, significant reduction of PI(3,4)P2 and enrichment of PI(3,4,5)P_3_ in knobs. Redistribution of the phospholipids accompanied marked elongation of cilia in INPP5E-OMP knockout (KO) OSNs. Such a dramatic remodeling of phospholipid composition however did not affect other integral membrane lipids (cholesterol, sphingomyelin, glycosylated phosphaditylinositol, phosphatidylserine). Proteins known to bind with high affinity PIP2 entered the cilia of the KO OSNs. Loss of INPP5E did not affect ciliary localization of endogenous olfactory receptor M71/M72 or distribution and movement of IFT122 particles implicating independent of phospholipids mechanism of retrograde protein transport in cilia of mature OSNs. Net odor sensitivity and response magnitude as measured by EOG was not affected by the mutation. However, odor adaptation in the KO mouse was significantly impaired resulting in less efficient recovery and altered inactivation kinetics of the odor response at the EOG and single-cell level. These findings implicate phosphoinositide-dependent regulation of active Ca^2+^ extrusion in OSNs whereby controlling the rate of sensory adaptation.

**Significance statement:** Currently there are little if any available treatment to cure congenital ciliopathies. This is in part due to lack of basic knowledge of cilia biology. Olfactory cilia as well as primary cilia appear to be a phospholipid privileged organelle distinct from the rest of plasma membrane albeit sharing its continuity. We characterized distribution of several critically important for cell biology phospholipids and showed that their balance, especially of PIP2, is disrupted in Joubert syndrome animal model and has functional implications. Virally assisted delivery of wild type *INPP5E* to the mutant OSNs was able to restore localization of PIP2 and rescued impaired response to odor.

## Introduction

Joubert syndrome (JS) is a recessive heterogeneous disorder heavily affecting structure and signaling mechanisms within the primary cilia (PC), an almost ubiquitous organelle on every eukaryotic cell. As a result JS has several severe conditions, a distinctive brain malformation “the molar tooth sign”, polycystic kidney, fibrosis liver and polydactyly (Valente et al., 2008). Most of the proteins encoded by genes involved in JS localize to a structure at the proximal part of PC called the transition zone (TZ) (Garcia-Gonzalo et al., 2011; Chih et al., 2012). The TZ anchors PC to the plasma membrane and regulates the movement of proteins in and out (Garcia et al., 2018). A few of the JS associated genes, among others ARL13B (Cantagrel et al., 2008), INPP5E (Jacoby et al., 2009), CSPP1 (Akizu et al., 2014) and IFT172 (Halbritter et al., 2013), encode proteins that localize to PC. Notably, ciliary localization of INPP5E depends on interaction with ARL13B, CEP164 and PDE6D comprising a network of the TZ proteins (Humbert et al., 2012; Nozaki et al., 2017). Therefore, INPP5E dysfunction due to the loss of catalytic activity and mislocalization is likely to play a key in the JS disease.

INPP5E gene encodes 5’-phosphatase converting polyphosphoinositide (PPI) lipids PI(4,5)P2 (PIP2), PI(3,5)P2 and PI(3,4,5)P3 (PIP3) into PI(4)P, PI(3)P and PI(3,4)P2, respectively (Kisseleva et al., 2000; Bielas et al., 2009; Hasegawa et al., 2016). Each of the lipids, representing a small fraction of all membrane associated lipids, plays indispensable role in regulating many aspects of cellular physiology including cell division, vesicle trafficking, control of transmembrane transport by utilizing specific adaptor proteins or direct interaction with effectors (Di Paolo and De Camilli, 2006; Balla, 2013). Recent studies have directly implicated INPP5E 5’-phosphatase and phosphoinositide kinase PIPKIγ in ciliogenesis arguing that interplay between PI(4)P and PIP2 may play a crucial role in organization of TZ hence controlling protein trafficking and signaling within cilium (Garcia-Gonzalo et al., 2015; Xu et al., 2016; Phua et al., 2017; Garcia et al., 2018). An overall emerging conclusion is that phospholipids are indispensable in controlling ciliary permeability and hence unique protein composition within this organelle (Verhey and Yang, 2016).

PIP2 and PIP3 have well established role controlling GPCRs, ion channels and transporters (Hilgemann et al., 2001; Brady et al., 2006; Suh and Hille, 2008; Hansen et al., 2011; Logothetis et al., 2015; Yen et al., 2018). PI(4)P has less defined role played in the plasma membrane except being a substrate for synthesis of PIP2 (Hammond et al., 2012) or providing important contribution to the overall electrostatic charge of the plasma membrane (Fairn and Grinstein, 2012). PI(3,5)P2 is thought to be exclusively present in endosomal membranes to control vesicle trafficking (Zolov et al., 2012), on contrary PI(3,4)P2 is generated at the plasma membrane and mediates endocytosis and focal adhesion (Marat and Haucke, 2016; Fukumoto et al., 2017). Foremost, PI(3,4)P2 as well as PIP2 and PI(4)P were directly implicated in maintenance of PC and GPCR trafficking through interaction with an adaptor protein TULP3 (Conduit et al., 2012; Mukhopadhyay et al., 2017; Garcia et al., 2018).

Our previous studies established multi-ciliated mouse olfactory sensory neurons (OSNs) as a unique system to study cell biology and functional role of cilia in developing and fully matured cells. Several ciliopathy-related genes, IFT88, BBS1 and BBS4 to be critically important for ciliogenesis and proper function of olfactory cilia (McIntyre et al., 2012; Williams et al., 2017; Green et al., 2018). Surprisingly, there is little knowledge about lipid composition of ciliary membrane of mammalian cells (Zhao et al., 2012; Lechtreck et al., 2013). Several studies suggested phosphoinositides, in particular PIP3, to modulate cyclic nucleotide-gated ion channel and inhibitory odor coding in mammalian OSNs (Brady et al., 2006; Ukhanov et al., 2011). Early evidence exists of odor-evoked hydrolysis of PIP2 (Boekhoff et al., 1990) substantiated by a more recent finding of translocation of PIP2 in the dendritic knob of mammalian OSNs (Klasen et al., 2010; Ukhanov et al., 2016). To systematically investigate role of PPIs in cell biology of olfactory system we generated an olfactory specific mutant mouse lacking functional *INPP5E* allowing for the first time to analyze distribution and functional implication of disease related change in ciliary phospholipid composition.

## Materials and Methods

### Animals

All procedures were approved by the University of Florida Institutional Animal Care and Use Committee, protocol 201608162. *INPP5E^flox/flox^* mouse has been made in the lab of Dr.Stephane Schurmans (University of Liege, Belgium). Mice were housed in a standard animal facility room at the University of Florida. To generate olfactory tissue specific mutant, INPP5Eflox/flox founders were crossed with OMP-Cre mouse (JAX stock#006668, deposited by Peter Mombaerts). Resulting INPP5E-OMP mice were genotyped using a standard PCR (Jacoby et al., 2009). Animals of both sexes were used in experiments.

### cDNA constructs and adenovirus production

Plasmids containing cDNA fragments were provided as follows (PLCδ1-PH-GFP, Addgene #51407, Btk-GFP Addgene #51463, mCherry-P4M-SidM, Addgene #51471, deposited by Tomas Balla; Tapp1-GFP, a gift from Takeshi Ijuin, Kobe University; D4H-mCherry, a gift from Gregory Fairn, University of Toronto; Lact-C2-GFP, Addgene 22852, deposited by Sergio Grinstein; Eqt2-SM-GFP, a gift from Christopher Burd, Yale University; TULP1 and TULP3, a gift from Saikat Mukhopadhyay, University of Texas Southwestern; Kir2.1, Addgene #32669, deposited by Matthew Nolan; PC2 (PKD2), Addgene #83451, deposited by Thomas Weimbs; Efhc1, a gift from Kazuhiro Yamakawa, RIKEN; IFT122, a gift from Jonathan Eggenschwiler, University of Georgia; GCaMP6f, Addgene 40755, deposited by Douglas Kim). C-terminal catalytic domain of INPP5E was subcloned from PJ-INPP5E (Addgene #38001, deposited by Robin Irvine). Full-length wild-type human INPP5E was cloned in the lab of Stephane Schurmans. MyrPalm lipid anchored constructs were described previously (Williams et al., 2014). All cDNAs were fused with GFP or mCherry and inserted into the pAd/CMV/V5-DEST^TM^ expression vector using Gateway technology (Invitrogen). Adenoviral vectors were propagated in HEK293 cells using the ViraPower protocol (Invitrogen), isolated with the Virapur Adenovirus mini purification Virakit (Virapur, San Diego, CA) and dialyzed in 2.5% glycerol, 25 mM NaCl and 20 mM Tris-HCl, pH 8.0 (Slide-A-Lyzer Dialysis Cassette, 10,000 MWCO) overnight. Alternatively, purified virus was dialyzed and further concentrated using ultrafiltration device Sartorius Vivaspin-6 (100,000 MWCO).

### Whole mount immunocytochemistry

Animals were sacrificed by inhalation carbon dioxide followed by cervical dislocation. Freshly dissected turbinates and septum were drop fixed for 3-4 hours on ice in freshly prepared 4% paraformaldehyde in a phosphate-buffered saline (PBS), pH7.4 supplemented with 20% sucrose. Tubes containing the tissue were carefully placed in refrigerator and left for the duration of fixation without any movement or agitation. This step was absolutely critical for the preservation of cilia known to be extremely sensitive to mechanical damage. The tissue was thoroughly washed in PBS and blocked with PBS containing 3% fetal bovine serum, 2% bovine serum albumin and 0.3% Triton X-100 for 2 h at RT. The tissue was then incubated with primary antibody against mouse M71/72 olfactory receptor (gift from Dr. Gilad Barnea, Brown University) raised in guinea pig, diluted 1:1,000 in the same blocking solution. Finally, the tissue was incubated with secondary anti-guinea pig Alexa 568 (1:1,000) for 2h and placed in antifading mounting agent Vectashield (Vector Labs) on the glass coverslip. Specimen were analyzed in the inverted confocal microscope Nikon TiE-PFS-A1R. Images were post-processed using Nikon Elements software (version 4.30) and NIH ImageJ (Wayne Rasband, NIH http://imagej.nih.gov/ij) and assembled in CorelDraw v.18 (Corel)

### En face imaging of adenovirally expressed proteins in live mouse OE

To express genes of interest, 10-20 µl of purified viral construct was intranasally administered to animals ranging 10-40 days of age. Typically, viral delivery was repeated in three consecutive days. Ten days post infection animals were anesthetized with CO_2_, rapidly decapitated, entire turbinates and septum were dissected and kept on ice in a petri dish filled with freshly oxygenated with carbogen modified artificial cerebrospinal fluid (ACSF) that contained (in mM): 120 NaCl, 25 NaHCO3, 3 KCl, 1.25 Na2HPO4, 1 MgSO4, 1.8 CaCl2, 15 glucose, 305 mOsm (adjusted with sucrose), pH7.4. For imaging a small piece of the OE was mounted in the perfusion chamber (RC-23, Warner Instruments) with apical surface facing down and analyzed in Nikon TiE-PFSA1R confocal microscope using preset configuration for acquisition of GFP and mCherry fluorescence.

For total internal reflection fluorescence microscopy (TIRF) *en face* imaging, virally transduced animals were prepared as above. TIRF imaging was performed on a Nikon Eclipse Ti-E/B inverted microscope equipped with a 100x oil immersion CFI APO TIRF 1.49 NA and an EMCCD camera (iXon X3 DU897, Andor Technology). Dedicated ImageJ plug-in was used to generate line-scan kymograms for measuring particle velocities from imported time series.

### Single cell GCaMP6f calcium imaging of odor evoked response

Calcium imaging was done as described previously (Ukhanov et al., 2016). Animals of 4-6 weeks of age were used for experiments 10-14 days post-administering adenovirus encoding GCaMP6f. Tissue was prepared and mounted the same way as described above. The chamber was transferred to the stage of upright microscope Zeiss Axioskop2F equipped with 40x 0.75NA water-immersion objective lens. Experimental solutions were applied directly to the field of view through a 100 µm diameter needle made of fused silica and connected to the 9-channel Teflon manifold. Each perfusion channel was controlled by the electronic valves (VC-6, Warner Instruments). The calcium response presented as an increase of GCaMP6f fluorescence emanating from the knob and underlying dendrite. The tissue was illuminated using a standard eGFP filter cube BP490nm/535nm (Omega Optical, USA) and the emitted light was collected at 530 nm (BP 530/20 nm, Omega Optical, USA) by a 12-bit cooled CCD camera (ORCA R2, Hamamatsu, Japan). Both the illumination system (Lambda DG-4, Sutter Instruments, USA) and image acquisition were controlled by Imaging Workbench 6 software (INDEC BioSystems). Before processing fluorescence intensity was corrected for the background. Each knob was assigned a region of interest (ROI) and changes in fluorescence intensity within each ROI were analyzed and expressed as the peak fractional change in fluorescent light intensity (F-Fo)/Fo where Fo is the baseline fluorescence before odorant application. A mixture of 132 distinct odorants were diluted from 0.5M DMSO stock and further diluted in ACSF to the working concentration. Odorants were delivered as aqueous solutions prepared in freshly oxygenated ACSF. A stock solution of a complex odorant mixture of 132 distinct chemicals (Ukhanov et al., 2013) was made in DMSO and then mixed to final 1:10,000 dilution with ACSF. ACSF supplemented with 0.1% DMSO, the odorant carrier, served as the control solution. All 132 odorous chemicals in the stock solution were at the same concentration of 3.79 mM.

Analysis and graphical presentation of calcium imaging data was performed with Imaging Workbench 6 (INDEC), Clampfit 10.7 (Molecular Devices), NIH ImageJ software, Microsoft Excel and Graph Pad Prism 6.

### Electroolfactogram recording

Mice were anesthetized with CO_2_, rapidly decapitated, and the head split along the cranial midline. Septal tissue was removed to expose olfactory turbinates. Vapor-phase odors were delivered by a pressurized nitrogen line connected to a sealed 100 ml glass bottle and directly injected into a continuous stream of humidified carbogen flowing over the tissue. Odorants were prepared by diluting pure stock into deionized water and final working concentration calculated as molar (v/v). Responses to odors were recorded with a standard glass micropipette tip-filled with agarose and backfilled with PBS using a Multiclamp 700A amplifier controlled by Multiclamp 700A and Clampex 9.2 software (Molecular Devices). EOG were measured as the maximal peak amplitude from the pre-pulse baseline using Clampfit 9.2 software (Molecular devices).

### Statistical Analysis

All statistical tests were done in Prism 6 (GraphPad) by using non-parametric Mann–Whitney test, unpaired t-test or one-way ANOVA, and *p* < 0.05 was considered to be statistically significant. All the group statistics are presented as mean ± SEM (standard error of mean).

## Results

### Loss of INPP5E causes PIP2 redistribution from the knob into cilia

To avoid neurodevelopmental deficiencies characteristic to the Joubert syndrome animals (Bielas et al., 2009; Jacoby et al., 2009) and restrict an effect of the INPP5E removal only to mature olfactory sensory neurons (OSNs) we followed established in our lab strategy of generating a conditional Cre recombinase-dependent mutant under the control of OMP promoter (Williams et al., 2014). INPP5E deficient mouse was generated by crossing INPP5E^flox/Δ^ (Jacoby et al., 2009) with *OMP-Cre* producing a conditional knockout INPP5E-OMP (KO). INPP5E hydrolyzes with high affinity two phosphoinositide species PI(4,5)P_2_ (PIP2) and PI(3,4,5)P_3_ (PIP3) generating PI4P and PI(3,4)P2, respectively (Kisseleva et al., 2000; Kong et al., 2000). Distribution of PIP2 in mature OSNs was measured using *en face* confocal microscopy of intact olfactory epithelium transduced with adenovirus encoding PLCδ1-PH (PLC-PH) domain tagged with GFP (Fig. 1). In WT littermate control mouse more than 50% of all infected with PLC-PH probe cells showed extremely polarized PIP2 distribution being very abundant in the OSN knob whereby labelling only a short ciliary appendages (Fig. 1A,C). Cilia otherwise remain intact and extend to its full length of 29.5±0.5 µm (n=753) revealed by labeling with an inert lipid probe MyrPalm fused to mCherry (Fig. 1A merged image, middle panel). In a small fraction of cells PLCPH probe decorated few relatively long appendages, probably spanning entire length of cilia summarized in Figure 1C. Overall distribution of PIP2 and MyrPalm-mCherry in the WT cilia result in a separate histograms summarized in Figure 1D. Loss of INPP5E function in the KO mouse severely impacts ciliary PIP2 distribution resulting in its homogenous redistribution along the entire length (Fig. 1B middle panel). Remarkably, this affected most if not every cilium (Fig.1C, KO) shifting distribution of PIP2 domain length to complete overlap with that of ciliary length marker MyrPalm-mCherry (Fig. 1E). Resulting mean cilia length in the mutant OSNs was significantly increased from 29.5±0.5µm in WT littermates to 35.3±0.6µm in the INPP5E-OMP KO mouse (n=495, unpaired t-test, t=7.363 df=1246, p<0.0001). Notably, abundance of PIP2 in the plasma membrane has not been changed as reported on the z-stack projection across the dendritic knobs (Fig. 1 WT, KO right most panels).

**Figure 1.**
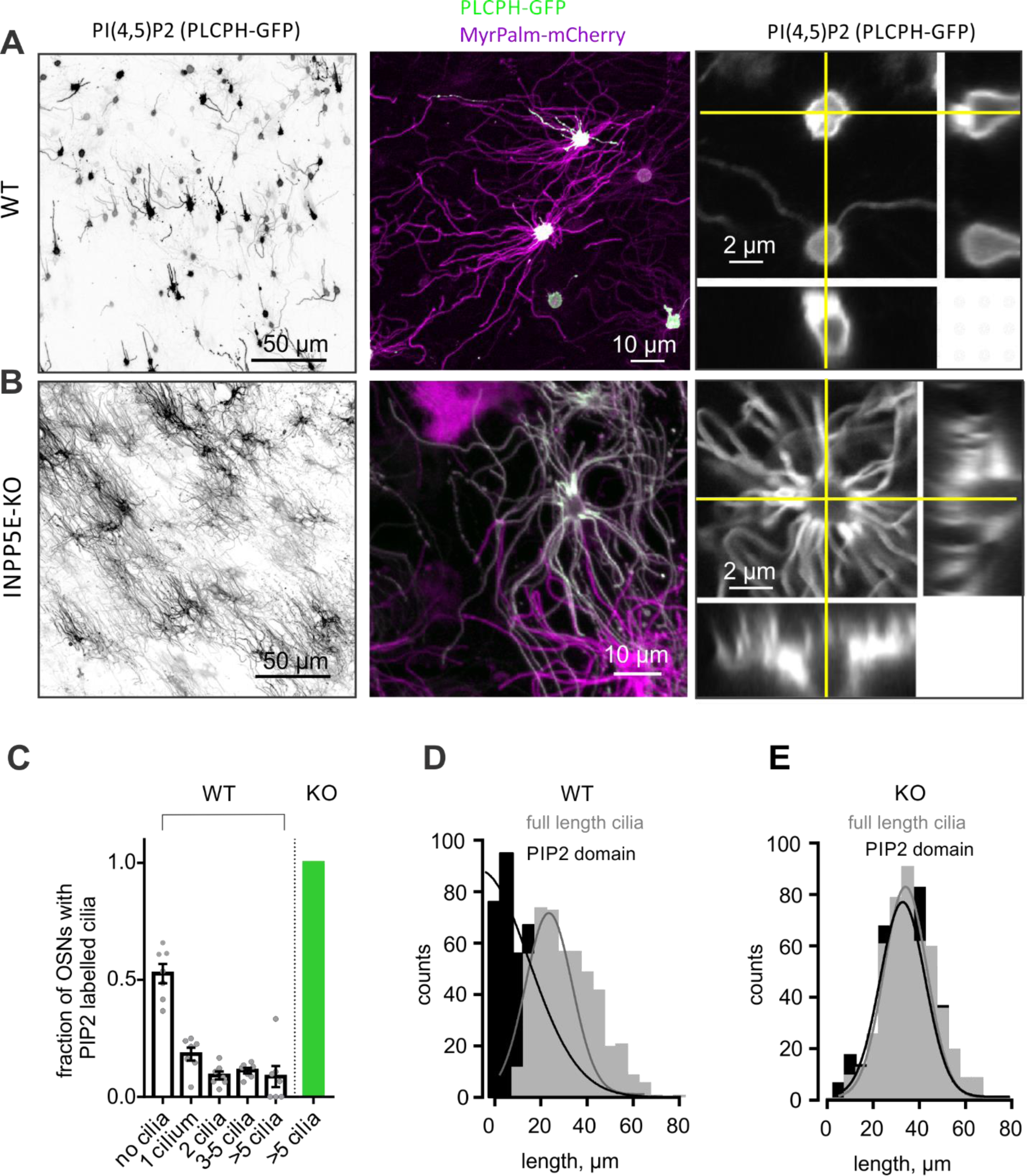
Loss of INPP5E causes redistribution of PIP2 and elongation of cilia in mouse OSNs. (A) PLCδ1-PH domain tagged with GFP (PLCPH-GFP), a probe for PIP2, was mostly restricted to the knob of the wild type (WT) OSNs. In a small percentage of OSNs proximal segment of varying length was also enriched in PIP2. Inert membrane bound lipid probe MyrPalm-mCherry was used as a counterstain to label the full length of axoneme opposing highly restricted localization of PLCPH-GFP resulting in overlapping colors (middle panel, white). PIP2 was evenly distributed in the plasma membrane of the knob as shown in z-stack view (right panel). Yellow lines denote z-stack projection shown at the bottom and right side of the image. (B) Opposite to the WT, in INPP5EOMP knockout PLCPH-GFP decorated the entire length of every cilium. This distribution resulted in a complete overlap of PLCPH-GFP and MP-mCherry labeling (middle panels, white color). The PIP2 redistribution is evident also in the z-stack view showing substantial enrichment of the proximal segment of cilia whereby PIP2 level in periciliary plasma membrane was not changed (right panel). (C) More than 50% of WT OSNs showed no PIP2 in their cilia. 18% of OSNs had only a single PIP2 positive cilium whereas three other groups of neurons equally represented remaining 30%. Conversely, PIP2 was detected in 100% of OSNs in INPP5E-OMP KO (green bar). (D) Length distribution of PLCPH-GFP positive aspects of cilia (PIP2 domain) in WT was substantially shifted to shorter values compared to the full cilia length measured with MyrPalm-mCherry, yielding 29.5±0.5 µm (n=753). (E) Distribution of both PLCPH-GFP and MyrPalm-mCherry length values showed complete overlap in INPP5E-OMP KO OSNs. Average ciliary length in the KO OSNs, 35.3±0.6 µm (n=495) was significantly longer than in the WT (unpaired t-test, t=7.363 df=1246, ****p<0.0001).

### Ectopic expression of wild type human INPP5E rescues PIP2 distribution in olfactory cilia of INPP5E-OMP OSNs

Rescue by viral gene delivery has been successfully accomplished in several mouse ciliopathy models (Beltran et al., 2012; McIntyre et al., 2012; Chávez et al., 2015; Green et al., 2018). Importantly, ectopic expression of wild-type INPP5E in mouse neuronal stem cells isolated from INPP5E knockout restored endogenous PIP2 and Grp161 levels in PC (Chávez et al., 2015). To address potential of virally assisted therapy of the Joubert syndrome ciliopathy model in vivo, we engineered a rescue adenoviral vector carrying a partial sequence of human *INPP5E* (NM_019892, residues 214-644) which was previously demonstrated to convert PIP2 in the plasma membrane of COS-7 cells (Hammond et al., 2012). CAAX prenylation domain was preserved at the C-terminus of INPP5E and conferred membrane anchoring. However, to our surprise catalytic domain of INPP5E alone was ineffective in converting PIP2 which was still redistributed along full-length cilia in INPP5E-OMP OSNs (not shown). Conversely, a full-length wild-type INPP5E (GFP-INPP5E-FL) was necessary and sufficient for rescue (Fig. 2). Ectopically expressed GFP-INPP5E-FL was enriched in the OSN knobs equally in WT (not shown) and the KO (Fig. 2A). GFP-INPP5E-FL localized to the entire length of cilia in INPP5E-OMP OSNs and dephosphorylated PIP2 resulting in a significant decrease of PIP2 ciliary domain length as measured with PLCPH-mCherry (Fig. 2B,C). Average length of PIP2 domain was in WT cilia 4.9±0.27 µm (n=110), in INPP5E-OMP cilia 28.5±1.37µm (n=54) and in rescued KO cilia 4.2±0.3µm (n=122), one-way ANOVA, F(DFn, DFd) 86.73 (2,283), p<0.0001.

**Figure 2.**
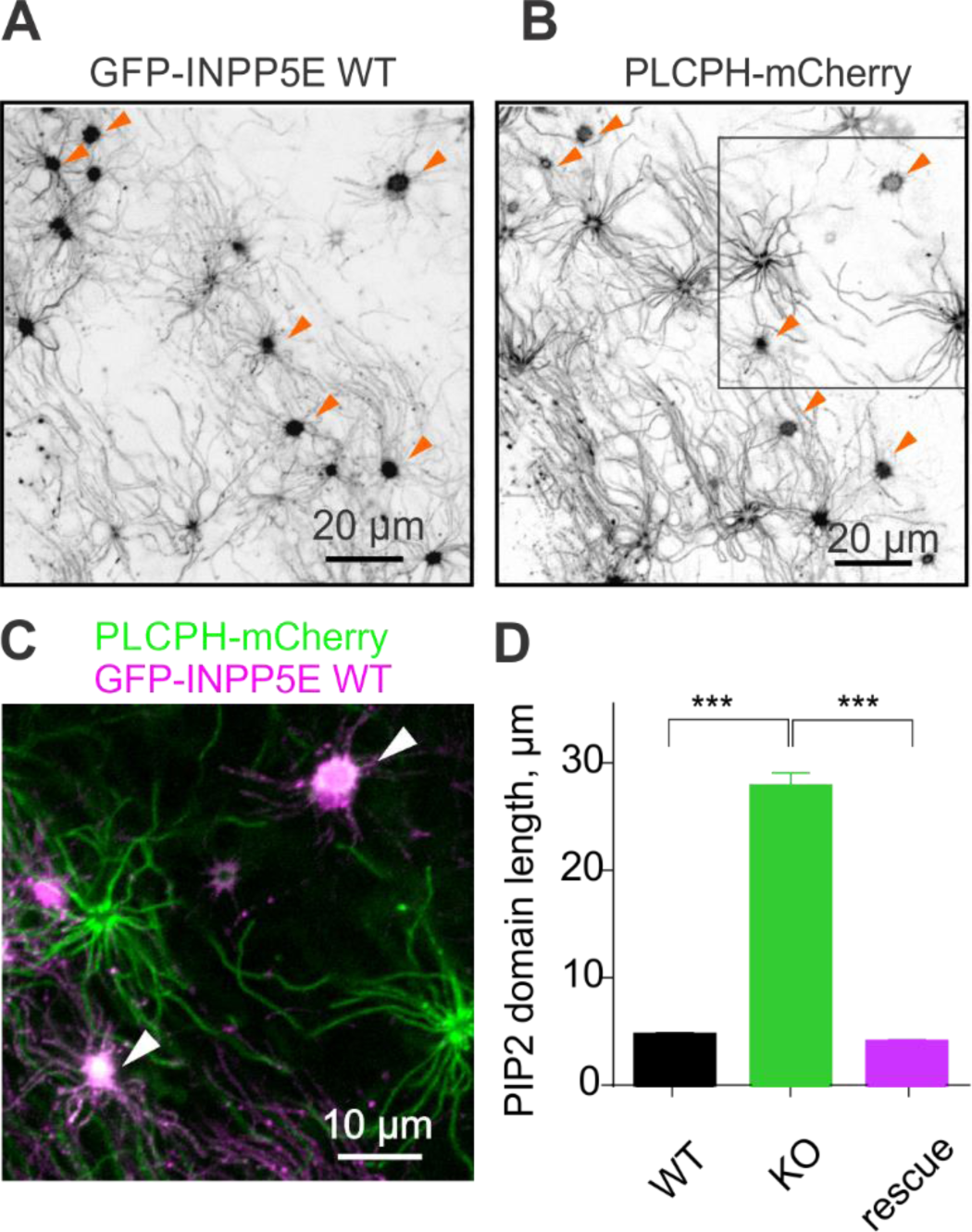
Virally induced ectopic expression of full-length wild type human INPP5E tagged with GFP (GFP-INPP5E-WT) completely reversed mislocalization of PIP2 in INPP5E-OMP KO mouse cilia. (A, B) INPP5E-OMP KO mice were infected at P8-P14 with a triple dose of Ad-GFP-INPP5EWT and tested 8-10 days later. GFP-INPP5E-WT is enriched in OSN knobs and also localized to cilia. The KO mice were co-infected with PLCPH-mCherry to measure rescue of the PIP2 localization. Several knobs of co-infected OSNs are indicated with arrowheads. (C) Zoomed-in view of the area marked with a square in (B) shows several knobs of OSNs co-infected with both viruses (arrowheads) resulting in a complete loss of ciliary PIP2 (green). (E) Rescue was quantified by measuring length of PIP2 positive ciliary aspect in the WT littermates and KO mice. The KO OSNs were identified within the same preparation by a strong ciliary distribution of PLCPH-mCherry, also lacking any detectable GFP-INPP5E-WT fluorescence. Rescue completely reversed INPP5E-OMP deficiency (cilia length 4.9±0.27 µm (n=110), WT; 28.5±1.37 µm (n=54), KO; 4.2±0.3µm (n=122), one-way ANOVA, F(DFn, DFd) 86.73 (2,283), ****p<0.0001).

### Deficiency of INPP5E also affects other phospholipids in OSNs

One of the main routes of PIP2 synthesis is thought to be by PI5K and PI4K kinase-dependent phosphorylation of PI(4)P and PI(5)P, respectively (Schramp et al., 2015). PI(4)P was shown to be highly enriched in PC of several cell types (Chávez et al., 2015; Garcia-Gonzalo et al., 2015) under the tight control of INPP5E which seems not to use PI5P as a substrate (Kisseleva et al., 2000; Madhivanan et al., 2015; Schramp et al., 2015; Conduit et al., 2017). A novel specific for PI4P probe, P4M-SidM tagged with mCherry showed surprisingly low abundance in the olfactory cilia of the control WT mice (Fig. 3A top panel). Conversely, in most of OSNs PI4P was highly enriched in the knob (Fig. 3A). We directly compared levels of PI(4)P in the knobs of WT and KO by measuring absolute fluorescence intensity. In the mutant OSNs mean level of PI4P showed slight but not significant decrease (Fig. 3D; 179±26 units, WT, n=94 and 143±17 units, KO, n=54; t-test, t=0.9777 df=146, p=0.3298). Besides PIP2, INPP5E also dephosphorylates PIP3 at even higher efficiency than PIP2, generating PI(3,4)P2 (Conduit et al., 2012). PI(3,4)P2 was measured by Tapp1-PH domain (Fukumoto et al., 2017) and found mostly restricted to the knobs with a minor presence in cilia and its overall distribution was not changed by the loss of INPP5E (Fig. 3B). However, quantitative analysis of PI(3,4)P2 revealed significant depletion in the knobs of INPP5E-OMP KO (Fig. 3E, 280±11 units, n=830, WT; 174±7, n=858; t-test, t=8.453 df=1686, p<0.0001). Finally, to assay distribution of PIP3 we used GFP tagged PH domain of Bruxton tyrosine kinase (Btk-GFP), a well characterized highly selective for PIP3 lipid probe (Balla, 2013). Again, PIP3 was highly enriched in OSN knob, showing relatively low penetrance into cilia similar to PI(4)P and PI(3,4)P2 (Fig. 3C upper panel). Several other selective for PIP3 probes based on the PH domains of ARNO, Akt and Grp1 proteins showed identical to Btk-GFP distribution in OSNs (data not shown). Intriguingly, quantitative analysis of PIP3 in OSN knobs of INPP5E-OMP KO showed dramatic increase by nearly 3-fold (Fig. 3F; 668±64 units, n=60, WT; 1,495±185, n=91, KO; unpaired t-test, t=3.536 df=149, p=0.0005) with very little if any build-up in cilia (Fig. 3C KO bottom panel). As a negative control we used inert membrane lipid anchor probe MyrPalm-mCherry which did not show any significant difference of the total abundance in OSN knobs in the WT and the KO (Fig. 3G, MP-mCherry, 340±31 units, n=46, WT; 378±23 units, n=70, KO; t-test, t=1.001 df=114, p=0.3188).

**Figure 3.**
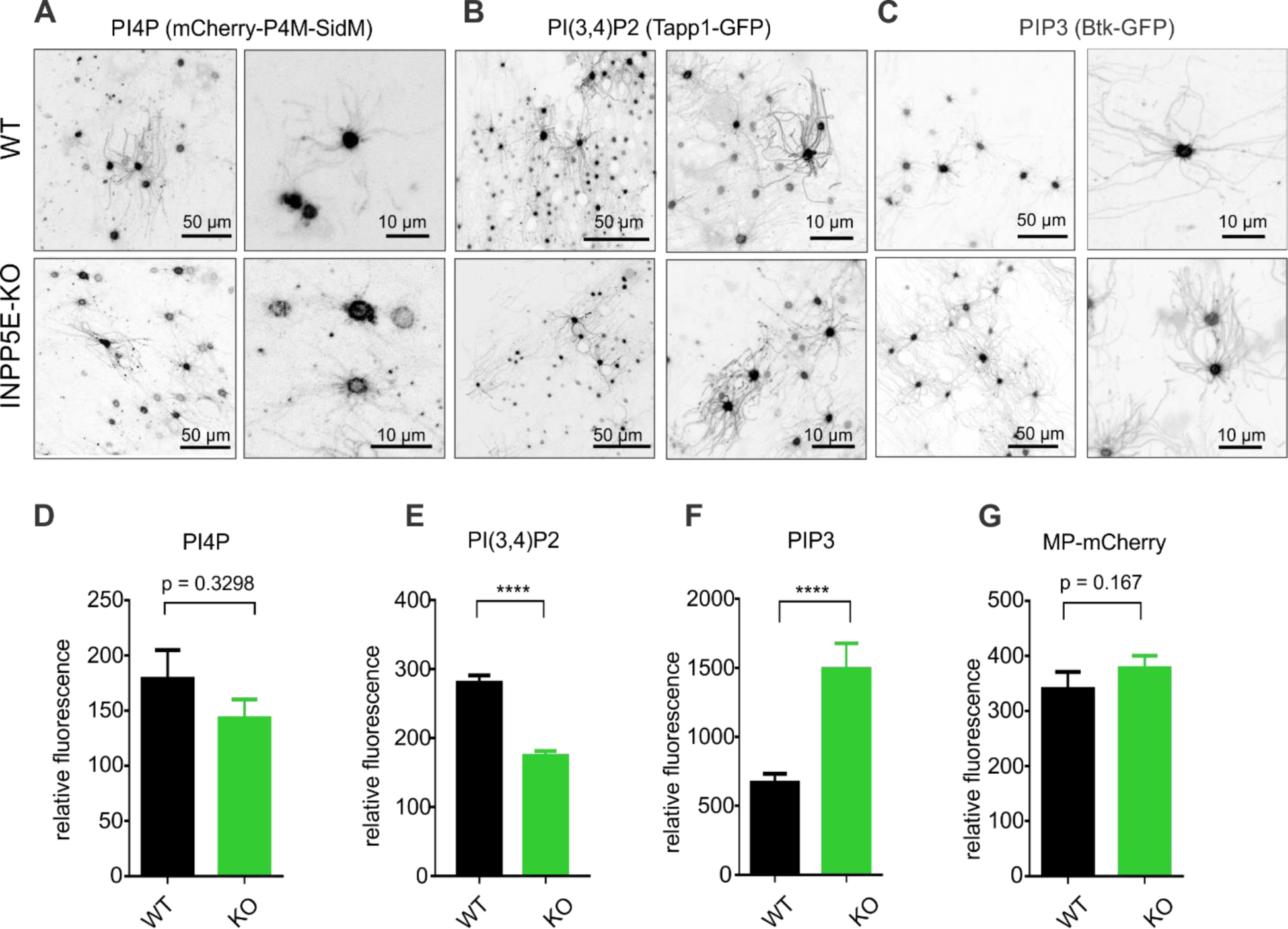
Other INPP5E-related phosphoinositides in mouse OSNs were almost exclusively restricted to the OSN knobs and change level in the INPP5E dependent manner. (A, D) PI(4)P probe, mCherry-P4M-SidM was not significantly affected by loss of INPP5E showing only insignificant trending decrease in the knobs (179±26 relative units, WT, n=94; 143±17 relative units, KO, n=54; unpaired t-test, t=0.9777 df=146, p=0.3298). (B, E) Tandem PH domain, Tapp1 tagged with GFP was used to specifically label membrane PI(3,4)P2 enriched mostly in the knobs and in cilia only in a small fraction of OSNs. Importantly, overall pattern of PI(3,4)P2 distribution did not change in INPP5E-OMP KO. Fluorescence intensity, however, measured in OSN knobs showed significant decrease in the KO compared to WT mice (280±11 relative units, n=830, WT; 174±7 relative units, n=858; unpaired t-test, t=8.453 df=1686, p<0.0001). (C, F) PIP3 detected with Btk-PH domain tagged with GFP, was restricted mostly to the knobs with relatively low presence in cilia of the WT and KO. Quantitative analysis of fluorescence showed increase of the intensity in the knobs of the KO (668±64 relative units, n=60, WT; 1,495±185 relative units, n=91, KO; unpaired t-test, t=3.536 df=149, ***p=0.0005). (G) Fluorescence intensity of MP-mCherry used as a negative control, was not significantly different in the OSN knobs of WT and KO mice (340±31 relative units, n=46, WT; 378±23 relative units, n=70, KO; unpaired t-test, t=1.001 df=114, p=0.3188).

### INPP5E does not affect overall lipid integrity of ciliary membrane

A growing consensus treats cellular plasma membrane as a patchwork of domains and PIP2 plays an important role regulating interaction of membrane with actin network and other components of membrane trafficking (Kwik et al., 2003; Stahelin et al., 2014). We hypothesized that the loss of INPP5E activity resulting in a substantial remodeling of ciliary PIP2 may impose additional collateral effect on overall lipid content. We first asked if cholesterol which is required for organizing membrane PIP2 rich domains may itself be reciprocally affected by its enrichment. D4H fragment of bacterial toxin perfringolysin-O recognizing cholesterol in inner membrane leaflet, tagged with mCherry (Maekawa and Fairn, 2015) selectively decorated proximal segment of cilia in WT mouse (Fig. 4A upper panel, arrowheads). Although D4H-mCherry was mainly associated with proximal zone it did label full length of cilia albeit not as strongly as MP-mCherry probe. Membrane enrichment in PIP2, however, did not affect overall localization of D4H-mCherry speaking to the lack of cross-talk between PPIs and cholesterol composition in olfactory cilia (Fig. 4A bottom panel).

**Figure 4.**
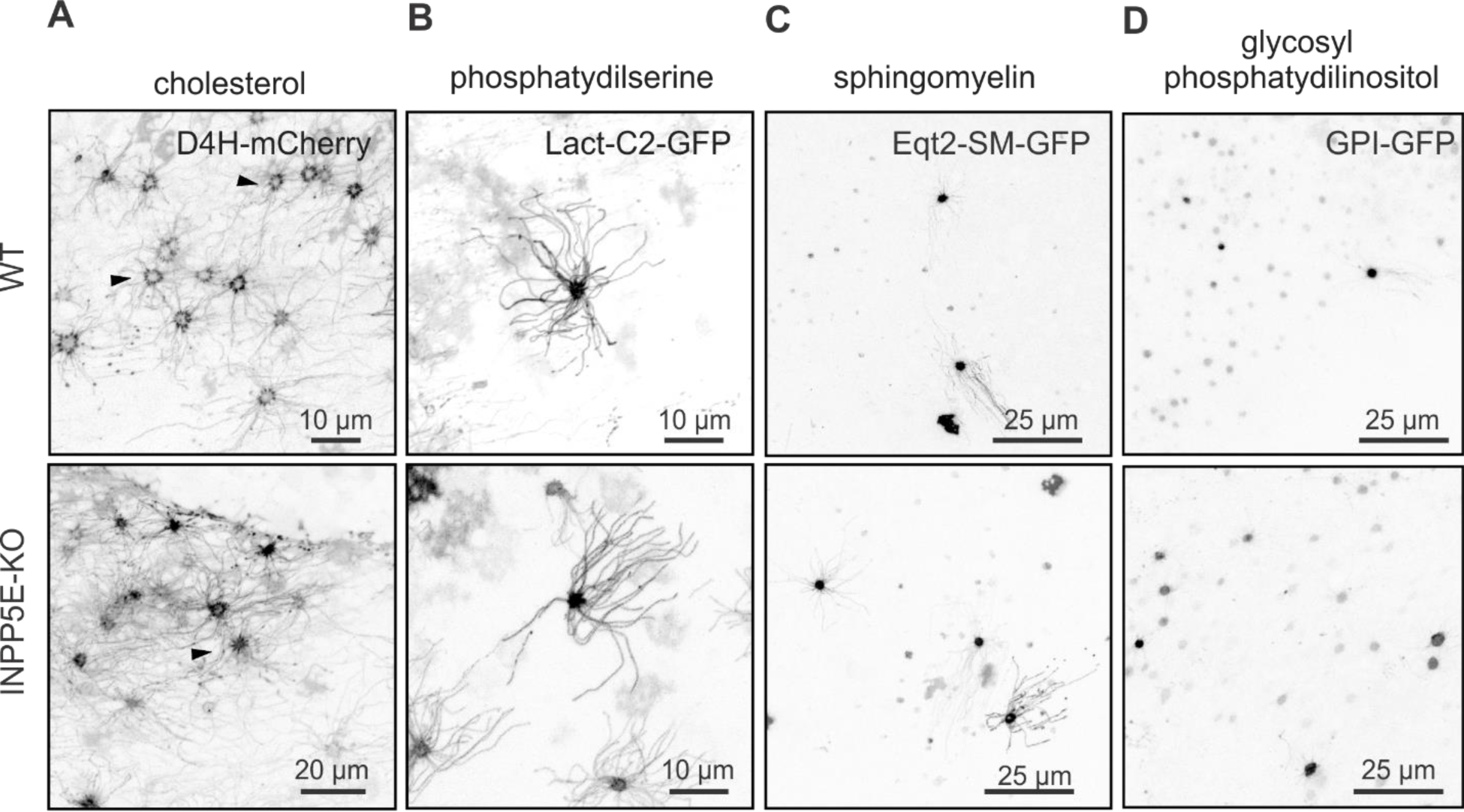
Distribution of integral membrane lipids was not changed in OSNs and cilia in INPP5E-OMP KO mice. (A) D4H-mCherry, a cholesterol binding probe was enriched in the proximal segment of olfactory cilia equally in the WT and KO OSNs (arrowheads). Cholesterol was detected albeit at a lower level also in the full-length ciliary axoneme. (B) Phosphatydilserine, probed with C2 motif of lactadherin, was uniformly distributed along the cilia and was also enriched in the dendritic knobs of OSNs. (C) Sphingomyelin specific probe, Eqt2-SMP-GFP was mostly enriched in the OSN knobs and detected at a low level in cilia. (D) Glycosylated phosphatydilinositol was probed in OSNs with a human folate 1 receptor, GPI-GFP which failed to detect any presence in cilia whereby mostly restricted to the knobs in both the WT and INPP5E-OMP KO mouse.

Another lipid required for the proper distribution of cholesterol is phosphatidylserine (Maekawa and Fairn, 2015). In accord with localization of cholesterol probe D4H-mCherry, ectopically expressed phosphatidylserine sensor Lact-C2-GFP, a fragment of lactadherin, was enriched in knobs and in addition evenly distributed along the entire length of cilia, and this pattern has not been affected in INPP5E-OMP mutant mouse (Fig. 4B).

We completed this screen by probing lipids relevant to protein trafficking and targeting, sphingomyelin and glycosylphosphatidylinositol (GPI). Eqt2-SM-GFP, equinatoxin-2 from the sea anemone *Actinia equina*, is a probe for sphingomyelin which is associated with Golgi-to-membrane vesicle trafficking (Deng et al., 2016). GPI-anchored proteins (e.g. human Folate Receptor-1) are directly targeted to the apical membrane in polarized cells, preferentially partitioning into cholesterol-rich raft domains (Paladino et al., 2008). Eqt2-SM-GFP showed partial enrichment of sphingomyelin in cilia of both WT and INPP5E-OMP KO mouse (Fig. 4C) whereby GPI-GFP was highly restricted only to the dendritic knobs in OSNs (Fig. 4D).

### Mutation in INPP5E impacts ciliary localization of proteins binding PIP2

Role of PIP2 and PIP3 has been long appreciated as regulators of multiple proteins thus forming structurally defined regions in the plasma membrane (Czech, 2000). Recently, several PIP2 binding proteins from a Tubby family implicated in ciliogenesis and ciliary protein traffic were shown to be mislocalized in PC of cells derived from INPP5E KO mouse (Mukhopadhyay et al., 2010). Members of Tubby-like protein family, TULP1 and TULP3 are anchored to the plasma membrane through its C-terminal PIP2 binding motif (Santagata et al., 2001; Mukhopadhyay and Jackson, 2011). We sought to visualize ciliary localization of TULP1 and TULP3 proteins not only to use a different PIP2 probe (Hammond and Balla, 2015) but also to test if translocation of proteins with affinity to PIP2 occurs in INPP5E-OMP deficient OSNs. Similar to PLCδ1-PH distribution, TULP1 and TULP3 were found mostly in the wild type OSN knobs (Fig.5A,B upper left panel). These findings support our earlier conclusion that PIP2 was excluded from cilia in the WT OSNs. However, in INPP5E-OMP mutant TULP1 and TULP3 translocated along the full-length axoneme, showing very similar to PLCδ1-PH distribution proving that PIP2 binding is sufficient for their ciliary entry (Fig. 5A,B bottom panels). Notably, TULP1 and TULP3 were particular enriched in the ciliary proximal segment of INPP5E-OMP KO (Fig. 5A,B right bottom panel, arrowheads). Overall, percentage of knobs showing TULP1-positive cilia was dramatically increased in the KO (Fig. 5A upper right panel, 25.49±0.06%, n=4, WT; 100%, n=6, KO; Mann-Whitney t-test, p=0.0048). Same distribution of TULP3 was found in cilia of the KO (Fig. 5B upper right panel, 30.87±0.12%, n=6, WT; 100%, n=4, KO; Mann-Whitney t-test, p=0.0095).

**Figure 5.**
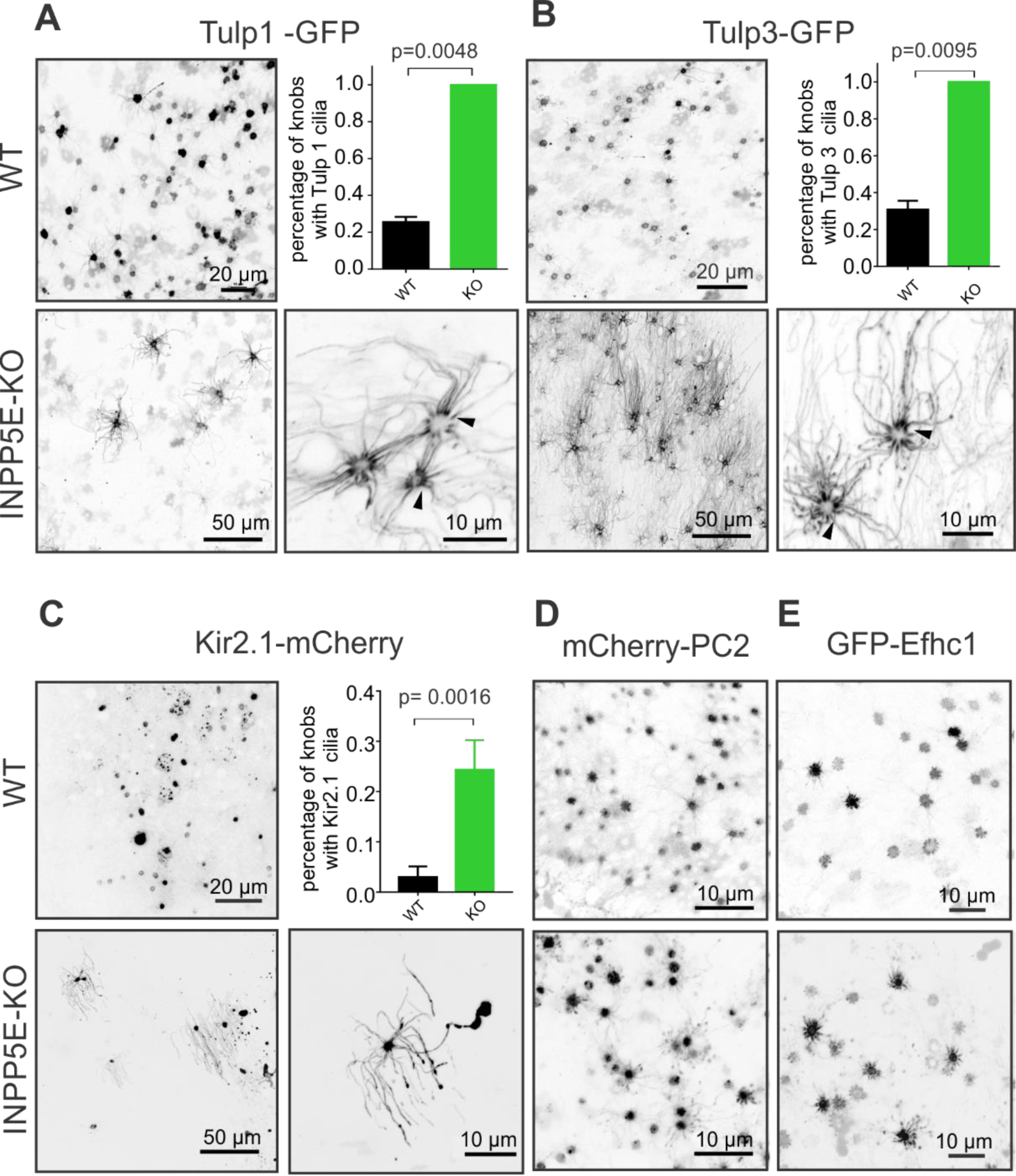
Soluble and polytopic proteins with affinity to PIP2 mislocalize in olfactory cilia of INPP5E-OMP KO. (A) Tubby-like proteins tagged with GFP, TULP1-GFP and TULP3-GFP were preferentially restricted to the knobs in the WT (upper left panel). Build-up of PIP2 in cilia of the KO resulted in complete redistribution of TULP1 (bottom panels). Note that loss of INPP5E activity led to a depletion of TULP1 within knobs revealing proximal segment of cilia decorated with TULP1-GFP (right bottom panel). Quantification of a percentage of the OSN knobs having TULP1-positive cilia per analyzed image showed significant increase in the KO (25.49±0.06%, n=4, WT; 100%, n=6, KO; Mann-Whitney t-test, p=0.0048). (B) TULP3-GFP, like TULP1, also showed dramatic redistribution between the knob and cilia resulting in a significant increase of percentage of knobs with TULP3-positive cilia (30.87±0.12%, n=6, WT; 100%, n=4, KO; Mann-Whitney t-test, p=0.0095). (C) Potassium inward rectifier ion channel, Kir2.1-mCherry, a polytopic protein with two membrane-spanning loops and a known affinity to PIP2 also changed its ciliary distribution in INPP5E-OMP OSNs. Kir2.1-mCherry moved into the ciliary membrane in a significantly larger fraction of OSNs in the KO (right upper panel, 3.02±0.02%, n=12, WT; 24.34±5.89%, n=10, KO; Mann-Whitney t-test, p=0.0023). (D, E) As a negative control we used a different ion channel, PC2 (PKD2 or TRPP1) tagged with mCherry (mCherry-PC2) and a microtubule binding protein Efhc1 (GFP-Efhc1) both of which, however, did not change their distribution in the knobs of INPP5E-OMP KO (upper panels, WT; bottom panels, KO).

Since TULP1 and TULP3 proteins endogenously localize to PC we asked if a different protein having more complex polytopic structure and known to bind PIP2 could be translocated into olfactory cilia in INPP5E-OMP mouse. Potassium inward rectifier channels (Kir1.x-6.x) and particular Kir2.x members that is endogenously expressed in olfactory system (Prüss et al., 2003) all depend on binding PIP2 for proper gating (Hilgemann et al., 2001; Hansen et al., 2011; Logothetis et al., 2015; Lee et al., 2016). Following this logic, we ectopically expressed in mouse OSNs Kir2.1-mCherry and found it highly localized to the knob with very little presence in cilia (Fig. 5C left panel). In INPP5E-OMP mouse Kir2.1-mCherry, however, moved into the ciliary membrane in a significantly larger fraction of OSNs (WT: 3.02±0.021%, n=12; KO: 24.34±5.89%, n=10; unpaired t-test, t=3.658, df=20, p=0.0016). Opposite to TULP proteins, other resident proteins expressed in PC (Efhc1, PC2) and not known to bind PIP2, failed to translocate into olfactory cilia of INPP5E-OMP mouse (Fig. 5D,E). Our findings support a role of PPI in mediating membrane delimited aberrant translocation of proteins in Joubert syndrome ciliopathy model.

### Loss of INPP5E function does not affect intraflagellar protein traffic and olfactory receptor localization

Recent evidence has shown that intraflagellar protein transport (IFT) is impaired in PC of several cell types, including embryonic fibroblasts and neuronal stem cells isolated from *INPP5E* KO mice (Mukhopadhyay et al., 2010; Chávez et al., 2015; Garcia-Gonzalo et al., 2015). Specifically, loss of INPP5E induces accumulation of IFT-associated proteins involved in anterograde and retrograde traffic which in turn disrupts proper retrieval of cargo proteins including relevant GPCRs (Chávez et al., 2015; Mukhopadhyay et al., 2017; Nozaki et al., 2017; Garcia et al., 2018). As such we envisioned a similar outcome in multi-ciliated OSNs of INPP5EOMP mutant mice. First, we ectopically expressed GFP-tagged IFT122 protein (IFT122-GFP) endogenously incorporated in IFT complex which forms particles moving along the olfactory cilia by kinesin and dynein motors (Williams et al., 2014). Velocity of intraciliary movement of IFT122-GFP was measured using TIRF microscopy in both WT and INPP5E-OMP KO and showed no effect of the mutation (Fig. 6A,B). Anterograde IFT velocity was 0.42±0.02 µm/s, n=336 in the WT; 0.42±0.01 µm/s, n=398 in the KO; unpaired t-test, t=0.1687 df=732, p=0.8661. Retrograde IFT velocity was 0.53±0.01 µm/s, n=360 in the WT; 0.54±0.01 µm/s, n=469 in the KO; unpaired t-test, t=0.545 df=827, p=0.5859. Second, ciliary localization of several endogenous GPCRs was perturbed in the PC in several INPP5E deficient models (Mukhopadhyay et al., 2010; Chávez et al., 2015; Park et al., 2015), hence we assayed endogenous distribution of olfactory receptor M71/72 using *en bloc* immunocytochemical approach. In WT and INPP5E-OMP KO we found very similar homogenous pattern of M71/72 localization along cilia (Fig. 6C,D). Cilia of 20-30 µm in length were observed both in WT and INPP5E-OMP KO which is on par with our standard method of live *en face* imaging (e.g. Fig.1 MyrPalm-mCherry).

**Figure 6.**
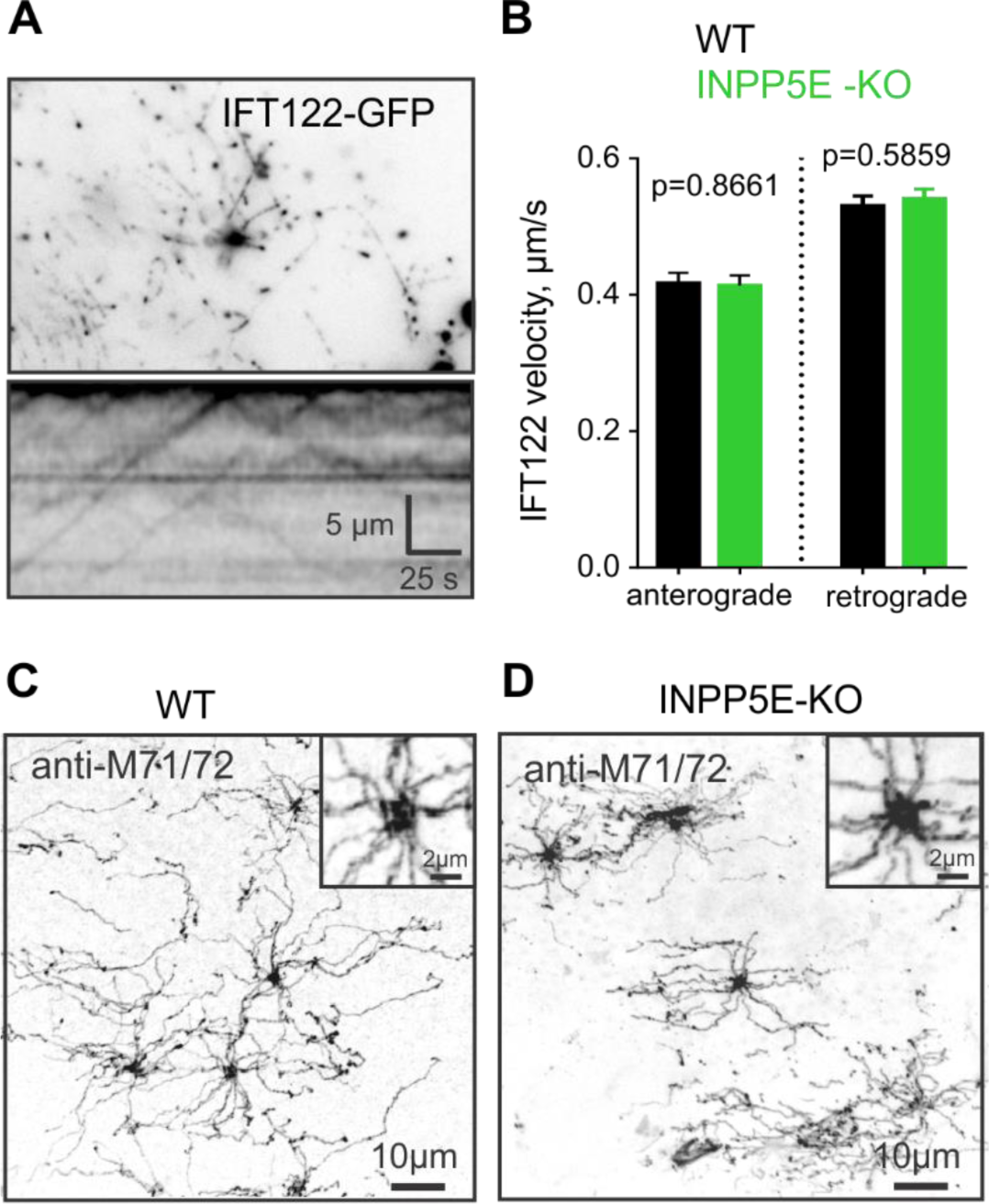
Velocity of intraflagellar transport (IFT) and endogenous localization of the olfactory receptor were not affected by loss of INPP5E function. (A) IFT particles incorporating ectopically expressed IFT122 protein, tagged with GFP (upper panel) was visualized by TIRF. Kymograph plot was generated by tracks of moving individual IFT particles (bottom panel). (B) Both the anterograde and retrograde transport of IFT122-GFP particles did not change due to the loss of INPP5E (anterograde IFT: 0.417±0.016 µm/s, n=336, WT; 0.414±0.015 µm/s, n=398, KO; unpaired t-test, t=0.1687 df=732, p=0.8661; retrograde IFT: 0.531±0.014 µm/s, n=360, WT; 0.542±0.014 µm/s, n=469, KO; unpaired t-test, t=0.545 df=827, p=0.5859). (C, D) En bloc immunolocalization in cilia of endogenous mouse olfactory receptor M71/72. Individual cilia often contained numerous particles due to the fixation artifacts. Also visible are fragments of punctated cilia detached during fixation. Otherwise intensity of labeling and overall distribution of M71/72, also shown in enlarged insets, was unchanged in the WT and INPP5E-OMP KO.

### Odor-mediated calcium transient is modulated by INPP5E

Based on the previously published role played by PPIs as modulators of ion channels including olfactory cyclic nucleotide-gated channel (Hilgemann et al., 2001; Brady et al., 2006), we hypothesized that unusually high steady-state accumulation of PIP2 in cilia as well as elevated PIP3 in the knob of INPP5E-OMP KO mouse may result in altered odor-evoked response. We measured odor-evoked calcium transients in the knob of OSNs ectopically expressing GCaMP6f (Fig. 7A left panel). Pressurized ACSF solution containing an odor mix made of 132 individual compounds representing several distinct chemical classes (Ukhanov et al., 2013) was applied to the OE surface through a glass micropipette whereby reproducibly activating many OSNs (Fig. 7A,B). A significant decrease of the time constant of termination phase of the GCaMP6f response was observed in INPP5E-OMP KO compared to the WT OSNs (Fig. 7C decay tau, 6,494±369 ms, n=167, WT; 3,596 ± 176 ms, n=110, KO, Mann-Whitney test, p<0.0001). Rise time from 10% to 90% of the GCaMP6f odor response amplitude was also significantly shorter in the KO (Fig. 7D, 1,118±71 ms, n=46, WT; 800±78 ms, n=30, KO, Mann-Whitney test, p<0.0001).

**Figure 7.**
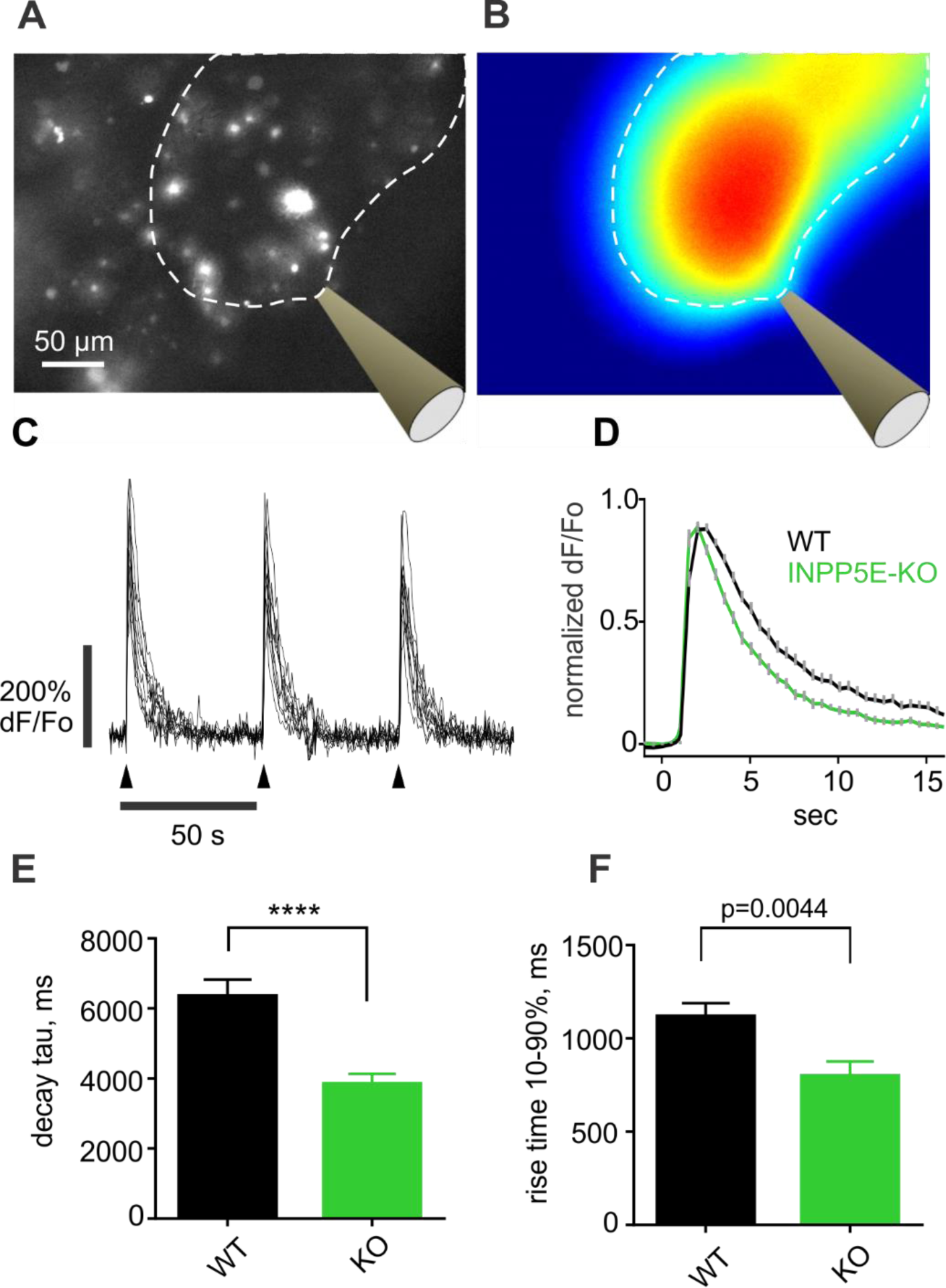
INPP5E is responsible for shaping the odor-evoked intracellular calcium transient in the knob of OSNs. (A, B) Ectopically expressed GCaMP6F was visualized in the *en face* preparation of mouse OE by wide-field fluorescence microscopy. Bright spots represent numerous OSN knobs. (B) Stimulation micropipette filled with a mixture of 132 different odorants diluted to 1:10,000 in ACSF was positioned as indicated. A single 100-ms pulse at 10psi pressure generated a plume of fluorescein covering an area over the epithelial surface demarcated by a dotted line. (C) Repetitive application of a single odor pulse (arrowheads) evoked nearly identical responses. GCaMP6F fluorescence corrected for background was calculated as (F-Fo)/Fo. (D) Individual traces measured in more than 100 OSNs across several areas and animals were averaged to create the graph. Traces were normalized to the peak value before averaging. (E, F) Odor-evoked GCaMP6F response had faster decay in INPP5E-OMP KO OSNs than the response in the WT control group (WT: 6.49±0.37 s, n=167; KO: 3.59±0.18 s, n=110, unpaired t-test, t=6.077, df=275, ****p<0.0001). The response in the KO also had faster rising phase (WT: 1.12±0.07 s, n=46; KO: 0.80±0.08 s, n=30, unpaired t-test, t=2.936 df=74, p=0.0044). To calculate termination phase time constant (decay tau) each individual trace was fit to an exponential function. Rise time 10-90% was defined as time to reach from 10% to 90% of the response peak level.

### Odor adaptation is impaired in INPP5E-OMP deficient mouse

Since calcium clearance from cilia and knobs of OSNs is critically involved in shaping the odor response (Stephan et al., 2012), we sought to further characterize odor response and odor sensitivity in INPP5E-OMP KO using electrophysiological approach. A short 100-ms pulse of amyl acetate of increasing concentration was applied to the tissue to build a concentration-response curve (Fig. 8A,B). Overall odor sensitivity was not changed in INPP5E-OMP (two-way ANOVA, F(5, 102) = 0.1858, P = 0.9674) resulting in overlapping dose-response (Fig. 8B). However, the kinetics of the response was different in the KO reminiscent of the changes observed in a single-cell GCaMP6F response. The EOG evoked by 10^-2^M amyl acetate reached its maximal magnitude faster (Fig. 8C; 10-90% rise time was 174.5±7.7 ms, n=37, WT; 157.9±10.9 ms, n=40, KO, unpaired Mann-Whitney test, p=0.0221). The response relaxed also faster to the baseline (Fig. 8D; termination phase was fit to a single exponential function yielding time constant of 4.57±0.15 s, n=81, WT; 3.40±0.16 s, n=28, KO; unpaired t-test, t=4.386, df=107, p<0.0001). Paired-pulse adaptation paradigm did not reveal any difference between the WT and KO using a short 100-ms pulse of amyl acetate (data not shown). However, we observed much stronger effect of the removal of INPP5E on adaptation of the EOG response to a longer 5-s pulse of 10^-3^M amyl acetate (Fig. 8E). Adaptation was measured as the ratio of the peak EOG evoked by the 2^nd^ odor pulse 40 s after the 1^st^ pulse (Fig. 8E,F black (WT) and green (KO) traces) and recovered slower in the KO (2^nd^/1^st^ peak ratio 0.733±0.026, n=18, WT; 0.514±0.022, n=9, KO; Mann-Whitney test, p<0.0001). The effect of the INPP5E-OMP deletion also resulted in a reduced plateau-to-peak ratio (ratio of 0.46±0.03 s, n=11, WT; 0.23±0.02, n=13, KO; Mann-Whitney test, p<0.0001). Finally, we analyzed decay kinetics by fitting termination phase of the EOG to a single exponential function yielding time constant of 1,707±124 ms, n=19, WT and 1,311±80 ms, n=20, KO (Mann-Whitney test, p=0.0083).

**Figure 8.**
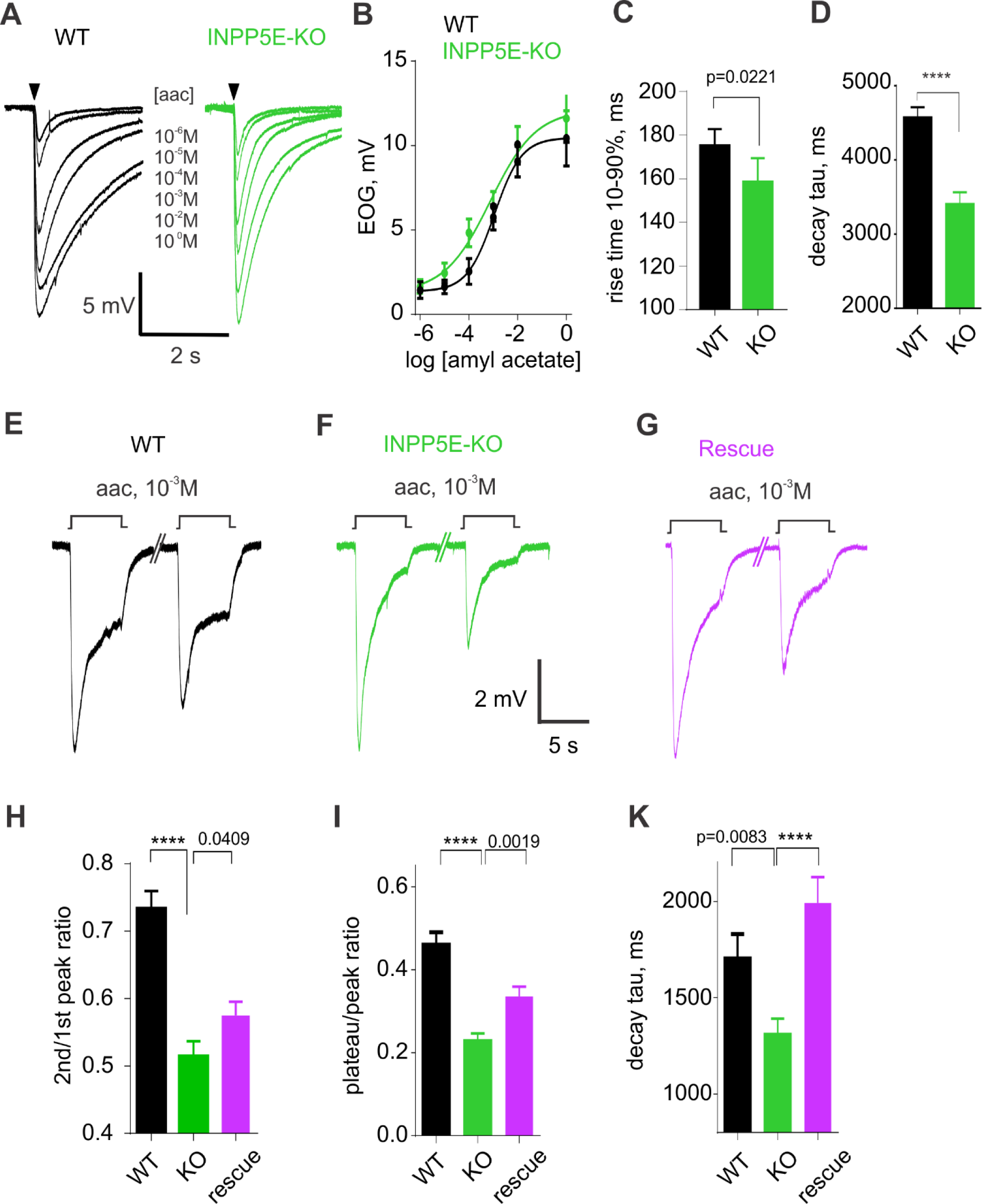
A faster single-cell odor response translates in more transient EOG in INPP5E-OMP KO. (A) Representative EOG traces recorded in response to 100-ms pulse of amyl acetate vapor, driven from the 90-ml head space of bottles containing increasing concentration ranging from 10^-6^M to a maximum of 1M (indicated at the individual traces). Odor application is denoted by a black arrowhead. (B) Dose-response relationship showed no significant difference between the WT and INPP5E-OMP KO (WT, n=7; KO, n=11; two-way ANOVA, F(5, 102) = 0.1858, P = 0.9674). (C, D) Rise time of the EOG evoked by a single 100-ms pulse of 10^-2^M amyl acetate (rise time 10-90%) was decreased in the KO compared to the WT (WT: 174.5±7.7 ms, n=37; KO: 157.9±10.9 ms, n=40; Mann-Whitney test, p=0.0221), similar to the time constant (decay tau) of the termination phase (WT: 4.57±0.15 s, n=81; KO: 3.40±0.16 s, n=28; unpaired t-test, t=4.386, df=107, ****p<0.0001). (E) EOG evoked by a longer 5-s pulse of 10^-3^M amyl acetate applied at the time indicated by a square step (aac, 10^-3^M) also appeared more transient in INPP5E-OMP KO (F). Ectopic expression of the full-length wild type INPP5E partially rescued the EOG shape (G). (H, I,K) Analysis of the ratio between peak amplitude of second and first EOG response, plateau-to-peak ratio and time constant of termination phase (decay tau) were significantly affected by the loss of INPP5E activity and restored by ectopic expression in OSNs of the wild type INPP5E. Second/first peak ratio (WT: 0.733±0.026, n=18; KO: 0.514±0.022, n=9; Rescue: 0.582±0.021, n=12; Mann-Whitney t-test, WT vs KO, ****p<0.0001; KO vs Rescue, p=0.0409). Peak/plateau ratio (WT: 0.462±0.028, n=11; KO: 0.230±0.017, n=13; Rescue: 0.336±0.024, n=16; Mann-Whitney t-test, WT vs KO, ****p<0.0001; KO vs Rescue, p=0.0021). Time constant of termination phase (WT: 1.707±0.124 s, n=19; KO: 1.311±0.080 s, n=20; Rescue: 1.991±0.134, n=16; Mann-Whitney t-test, WT vs KO, p=0.0083; KO vs Rescue, ****p<0.0001).

To assay feasibility of functional rescue, EOG was measured in adult INPP5E-OMP KO mice after 10 days of adenoviral treatment with Ad-GFP-INPP5E-FL. A 5-s pulse of 10^-3^M amyl acetate in the virally treated animals evoked odor response showing prominent recovery of the adaptation and kinetics (Fig. 8G-K magenta trace, Rescue). Statistical analysis confirmed a significant change of the EOG parameters (Fig. 8H-K). Ratio between 2^nd^/1^st^ EOG peak amplitude was increased following rescue treatment (0.514±0.022 n=9, KO; 0.582±0.021, n=12, Rescue; Mann-Whitney test, p=0.0409; one-way ANOVA comparing WT, KO and Rescue groups, F (2, 36) = 19.85, p<0.0001). Plateau-to-peak ratio was also increased (0.230±0.017, n=13, KO; 0.336±0.024, n=16, Rescue; Mann-Whitney test, p=0.0019; one-way ANOVA comparing WT, KO and Rescue groups, F (2, 37) = 21.99, p<0.0001). The decay time constant increased in rescued relative to the KO animals (KO, 1.31±0.08 s, n=20, KO; 1.99±0.13 s, n=16, Rescue; Mann-Whitney test, P<0.0001; one-way ANOVA comparing WT, KO and Rescue groups, F (2, 52) = 9.134, p=0.0004).

Our findings provide a compelling evidence of the role of phosphoinositides as a modulator of the odor response and their implication in ciliary biology of native multi-ciliated olfactory sensory neurons. The ability to rescue ciliary PIP2 distribution and the whole tissue odor response speak to the applicability of virus gene therapy treatment of Joubert syndrome related phenotype.

## Discussion

In the current study we sought to systematically study phospholipid content in olfactory cilia of the ciliopathy mouse model of Joubert syndrome. Conditional olfactory mutant has been generated by crossing INPP5EΔ/loxP and OMP-Cre mice to produce INPP5E-OMP knockout. We showed that loss of INPP5E activity in mature OSNs no longer restricts PIP2 at the TZ of cilia and adjacent knob membrane, changes abundance of PI(4)P, PI(3,4)P2 and PIP3 in the knob. Such dramatic phospholipid remodeling elongated cilia and facilitated the rate of intracellular calcium clearance in the OSN knob whereby affecting long term adaptation of the odor response.

Great effort has been put over the past two decades in deciphering the role of PPIs in different compartments of cell. Paradoxically, very little information is available on the ciliary membrane composition except for a few studies of the PC in *Chlamydomonas* and an actin-based hair cell stereocilia (Zhao et al., 2012; Lechtreck et al., 2013). Multi-ciliated mammalian OSNs is an attractive model to study not only basic cell biology of cilia but also its function as a specialized chemosensory antenna by analogy to the PC. Due to its substantially longer than PC length and unique exposure to external environment, olfactory cilia can be easily studied with imaging techniques in its native environment. Early published work revealed similarities between olfactory cilia and PC including molecular identity of components mediating trafficking and maintenance (McEwen et al., 2008; McIntyre et al., 2012; Williams et al., 2014).

Our results on the remodeling of PIP2 in cilia well correlate with previously published studies of INPP5E role in PC in different cell types of mammalian, fish and insect origin (Chávez et al., 2015; Garcia-Gonzalo et al., 2015; Park et al., 2015; Xu et al., 2017). However, there are several distinct differences placing olfactory cilia in its own class. In WT OSN cilia we did not confirm PI(4)P enrichment that could be decreased in INPP5E deficient cilia. The *INPP5E* gene (human ortholog *INPP5E*) is conserved in animal kingdom having some similarity even with protist green algae *Chlamydomonas reinhardtii*. Preferentially, INPP5E class phosphatases act on membrane-bound PPIs and to some extent targeting soluble species (Pirruccello and De Camilli, 2012; Pirruccello et al., 2014). Since WT knobs are enriched with PI(3,4)P2 it may hypothetically serve as a product of PIP3 hydrolysis by INPP5E or phosphorylation substrate for PI5K, a process typical for endosomal membranes (Posor et al., 2013). INPP5E in the medulloblastoma PC is mostly involved in converting PIP3 into PI(3,4)P2 (Eramo and Mitchell, 2016). However, the canonical route of PIP3 synthesis is via recruiting of class I and II PI3K occurring mostly during tyrosine kinase receptor activation during proliferation (Hawkins and Stephens, 2016). In OSNs recent studies showed that PI3K has low basal activity hence maintaining low membrane pool of PIP3 and only transiently activated following odor exposure (Ukhanov et al., 2010). Paradoxically, we found that steady-state ciliary distribution of PPIs other than PIP2 was not significantly changed. This suggests that due to redundancy between multiple 5’-phosphatases, resting PPIs level may not be at all strongly affected. In other study it took a combined knock-down of several isoforms of 5’-phosphatase, SHIP1, SYNJ1/2, OCRL, and INPP5EB to reveal significant elevation of PIP3 and still only a slight decrease of PI(3,4)P2 (Malek et al., 2017).

Published so far evidence strongly supports ciliary PI(4)P being a product of INPP5E-dependent hydrolysis of PIP2 and keeping the latter in the TZ domain. PI(4)P plays a role in regulation of endosome trafficking and has much less defined role in the plasma membrane except being a substrate for synthesis of PIP2 (Hammond et al., 2012) or providing important contribution to the overall electrostatic charge of the plasma membrane (Fairn and Grinstein, 2012). Importantly, PI(4)P and PIP2 comprise two nearly independent pools in the membrane and previous studies showed that PI(4)P may exist in the membrane for a short time as intermediate product channeled from PI4K to PI5K kinases to make PIP2 (Fairn and Grinstein, 2012). Because of this very dynamic process a steady-state level of PI4P may stay at nearly undetectable low concentration which may occur in OSNs.

One important consideration comparing PC and olfactory cilia in mature OSNs is that ciliogenesis of the PC is typically induced in cell culture by serum starvation or during embryogenesis by growth factors targeting Shh and/or tyrosine kinase receptor PI3K/PIP3/Akt pathways (Eramo and Mitchell, 2016). Probably, at the terminal developmental stage OSNs lack input from Hedgehog signaling negatively regulated by PIP2 in the PC through TULP-3 and Gpr161 (Mukhopadhyay et al., 2010; Chávez et al., 2015; Wrighton, 2015; Xu et al., 2016). Noteworthy, ciliary localization of INPP5E is heavily regulated by Arl13b, PDE6δ and MKS1 as a ciliary gate keepers controlling entry of lipidated cargo (Humbert et al., 2012; Fansa et al., 2016; Slaats et al., 2016; Nozaki et al., 2017). Congruently, MKS-5 (mammalian ortholog Rpgrip1L), a protein in control of the TZ assembly in *C.elegans*, was shown to control periciliary localization of PIP2 (Jensen and Leroux, 2017). Not surprisingly, very recent study has directly shown the role of PIP2 regulating integrity of TZ during ciliogenesis in Drosophila male germline (Gupta et al., 2018). Overall there is an emerging concept of the TZ as a lipid gate for ciliary protein traffic (Reiter and Leroux, 2017). IFT-A retrograde ciliary transport of Grp161 receptor involves IFT-122 protein binding TULP3 which is in turn recruited to the membrane by PIP2 (Mukhopadhyay et al., 2010; Boubakri et al., 2016). This mechanism is also responsible for proper ciliary trafficking of mechanosensitive ion channels NompC and PKD2 in *Drosophila* and *C.elegans*, respectively (Bae et al., 2009; Mukhopadhyay et al., 2010; Park et al., 2015). BBSome core proteins directly involved in IFT through interaction with kinesin and dynein motors are able to bind *in vitro* several PPIs with highest affinity to PI(3,4)P2 (Jin et al., 2010). All together with the role of INPP5E- and PIP2-dependent mechanism of IFT-A in the PC it is increasingly clear that PPIs comprise an important regulator of cilia function. To this end we sought to connect INPP5E function to any of the deficits in protein localization and trafficking. Surprisingly, we did not find any abnormality in ciliary localization of olfactory receptor M71/72 or in the velocity of IFT-A dependent transport of IFT122 protein. It is entirely possible that by the time *OMP* gene becomes active at E15 in embryogenesis INPP5E-releated processes in young OSNs are already established. Therefore we may have to look in even earlier developmental time point to determine an effect of INPP5E known to be indispensable for normal embryogenesis (Bielas et al., 2009; Jacoby et al., 2009).

Mislocalization and ensuing disfunction of INPP5E in mutants of Arl13b and CEP290, provided compelling evidence of its role in cilia elongation (Lu et al., 2015; Srivastava et al., 2017). Paradoxically, however in RPE1 cell line removal of Arl13b also mislocalized INPP5E but instead shortened the PC (Nozaki et al., 2017) suggesting more complex phenotype downstream of Arl13b. We showed that cilia elongation in OSNs of Joubert model validates importance of INPP5E to the cilia maintenance. However, as pointed earlier redistribution of PIP2 may induce aberrant remodeling of ciliary core by recruiting F-actin filaments thus contributing to cilia elongation (Madhivanan et al., 2015; Phua et al., 2017; Drummond et al., 2018). Ciliopathy associated with altered PIP2 distribution is not limited only by Joubert (INPP5E) syndrome but also occurs in a similar disease of oculo-cerebro-renal syndrome of Lowe (OCRL). OCRL (INPP5EF) is also a PPI 5’-phosphatase localized to the PC and its deficiency manifest severely impaired ciliogenesis (Madhivanan et al., 2015; Aguilar et al., 2016). As in case of INPP5E in mouse OSNs, loss of OCRL function leads to elongation of PC in renal epithelial cells (Rbaibi et al., 2012). If it holds true and OCRL localizes in OSNs along with INPP5E it may add even tighter control of the PIP2 in cilia. Probably it is logical to assume that OCRL acts on a distinct PIP2 pool either in the dendritic knob of OSNs without interfering INPP5E. Otherwise in INPP5E-OMP mutant ciliary PIP2 distribution was not changed.

Since PIP2 and PIP3 have been implicated in regulation of vast array of proteins including ion channels and transporters (Hille et al., 2015; Hilgemann et al., 2018), we hypothesized that redistribution of PIP2 in OSN cilia of INPP5E-OMP KO would affect its ability to transduce odor. Surprisingly, we found only a weak phenotype resulting in *acceleration* of odor response. Indeed, a number of proteins not immediately involved in signal transduction, like OMP, Goofy and most recently discovered Cfap69 were associated in a loss-of-function mutants with altered odor sensitivity (Buiakova et al., 1996; Kaneko-Goto et al., 2013; Talaga et al., 2017). It is therefore tempting to speculate that these and probably other orphan house-keeping proteins comprising olfactory cilia proteome (Klimmeck et al., 2008; Mayer et al., 2009) regulate ciliogenesis and its sensory function. Goofy protein is abundantly expressed in Golgi and likely regulates protein trafficking, in particular AC3 from Golgi to cilia and axons (Kaneko-Goto et al., 2013). Ciliogenesis in *Goofy* knockout OSNs was impaired yielding shortened cilia reminiscent of other ciliopathies (Kaneko-Goto et al., 2013). However, role of OMP has not been well understood likely affecting Na-Ca exchange and cAMP kinetics hence affecting overall odor response (Kwon et al., 2009; Dibattista and Reisert, 2016). Mutation of INPP5E in contrast to OMP, facilitated the rate of calcium extrusion from knobs and probably cilia well supported by the role of PIP2 as a positive regulator of cardiac Na-Ca exchanger (He et al., 2000). However, PIP2 build-up in cilia did not cause any adverse effect on overall signaling affecting odor sensitivity, maximal odor evoked EOG or short-term adaptation. Instead, it impaired longer form of adaptation induced by a long 5-s odor pulses. Our finding parallels an earlier evidence of the role of CaMKII kinase that controls long form of adaptation opposite to the short form (Leinders-Zufall et al., 1999). CaMKII is not known to be modulated by PPIs, however, there is another phosphoinositide-dependent kinase 1 (PDK1) known to activate several ACG kinases including protein kinase A both found in the PC and olfactory cilia whereby modulating signal transduction and adaptation (Cheng et al., 1998; Mashukova et al., 2006; Mayer et al., 2009; Mick et al., 2015).

We may hypothesize that simple leakage of PIP2 from the knob membrane into cilia may not be sufficient to re-establish lipid subdomains which thought to be crucial in organizing ion channels and other signaling molecules near the membrane (Hammond, 2016). Given that other INPP5E class phosphatases co-exist in OSNs along with INPP5E (Kanageswaran et al., 2015), it would be imperative in future studies to address growing complexity of the PPI pathways in ciliogenesis and ciliary signaling in olfactory system through its development. This complexity may probably explain why we could only partially rescue INPP5E-OMP KO phenotype. Viral delivery of wild-type INPP5E was sufficient to completely restore native distribution of PIP2 and partially recover odor response.

In conclusion, our work provides a novel insight in the lipid composition of olfactory sensory neurons and cilia in normal and disease-related condition. Having several commonalities to the INPP5E related phenotype described in the PC from different cell types, we observed distinct features placing multi-ciliated sensory neurons in its own class.

## Footnotes

This work was supported by US Public Health Service Grant NIH/NIDCD R01 - DC - 009606 (JRM), University of Liege (SS). The authors declare no competing financial interests.

